# A cortical microcircuit model reveals distinct inhibitory mechanisms of network oscillations and stability

**DOI:** 10.1101/2025.02.23.639719

**Authors:** Farzin Tahvili, Martin Vinck, Matteo di Volo

## Abstract

We identify a computational mechanism for network oscillations distinct from classic excitatory-inhibitory networks – CAMINOS (Canonical Microcircuit Network Oscillations) – in which different inhibitory-interneuron classes make distinct causal contributions to network oscillations and stability. A computational network model of the canonical microcircuit consisting of SOM, PV and excitatory neurons reproduced key experimental findings, including: Stochastic gamma oscillations with drive-dependent frequency; precise phase-locking of PV interneurons and delayed firing of SOM interneurons; and the distinct effects of optogenetic perturbations of SOM and PV cells. In CAMINOS, the generation of network oscillations depends on *both* the precise spike timing of SOM and PV interneurons, with PV cells regulating oscillation frequency and network stability, and delayed SOM firing controlling the oscillation amplitude. The asymmetric PV-SOM connectivity is found to be the key source ingredient to generate these oscillations, that naturally establishes distinct PV and SOM-cell spike timing. The CAMINOS model predicts that increased SOM/PV densities along the cortical hierarchy leads to decreased oscillation frequencies (from gamma to alpha/beta) and increased seizure susceptibility, suggesting a unified circuit model for oscillations across different frequency bands.

## Introduction

Neocortical microcircuits have a stereotypical organization and consist of multiple classes of excitatory and inhibitory neurons with well-defined morphological, electrophysiological and genetic characteristics (Rudy et al., 2011; Batista-Brito et al., 2018; Pfeffer et al., 2013; Markram et al., 2004; Douglas and Martin, 2004; Binzegger et al., 2004). The reciprocal interactions between these cell types are fundamental for stabilizing network interactions and shaping the nature of cortical dynamics, ranging from asynchronous to rhythmic activity in distinct frequency bands (van Vreeswijk and Sompolinsky, 1996; Buzsáki, 2006; Wang, 2010). The way in which inhibitory feedback controls stability and network dynamics in classic two-population E/I models (Wilson-Cowan, PING; Pyramidal-Interneuron-Network-Gamma models) is relatively well understood (Wilson and Cowan, 1972; Kopell et al., 2000; Wallace et al., 2011; Buzsáki and Wang, 2012). Yet, an understanding needs to be reached of the fundamental properties of networks containing multiple inhibitory classes, which may potentially play distinct roles in controlling network stability and dynamics.

In the activated state, e.g. during sensory stimulation, cortical circuits often engage in 20-80 Hz network oscillations, which are thought to play important functions in sensory and cognitive processing and to be implicated in various neurological and psychiatric disorders (Vinck et al., 2023; Singer, 1999; Fries, 2015; Cardin et al., 2009; Uhlhaas et al., 2008; Ray and Maunsell, 2015; Buzsáki and Draguhn, 2004; Fernández-Ruiz et al., 2021; Colgin et al., 2009). Several studies used optogenetics or targeted electrophysiological recordings to examine the distinct roles of PV and SOM interneurons in the generation of neocortical gamma oscillations. Both Veit et al. (2017a) and Chen et al. (2017) used optogenetic in mouse V1 to show that silencing SOM interneurons had a robust effect of reducing the power gamma oscillations. Yet, Chen et al. (2017) found that PV silencing reduced gamma-band oscillations while increasing low-frequency oscillations, while Veit et al. (2017a) found that PV silencing did not abolish gamma oscillations, although very strong PV suppression destabilized network dynamics. Onorato et al. (2023) recorded from SOM and PV interneurons and showed that PV interneurons show much stronger phase-locking than SOM interneurons and broad-waveform interneurons, consistent with several previous studies showing showing prominent gamma phase-locking in fast-spiking interneurons as compared to broad-waveform neurons (Hasenstaub et al., 2005; Csicsvari et al., 2003; Vinck et al., 2013; Perrenoud et al., 2016; Vinck and Bosman, 2016). Onorato et al. (2023) further show a substantially advanced phase relative to SOM interneurons, which may suggests that these two GABAergic interneuron populations control the timing of excitatory neurons at distinct phases. Altogether, these diverse empirical studies present a set of findings that are hard to reconcile with each other and are consistent with different theoretical models (Figure 1A): (1) An E-PV based synchronization mechanism, which is assumed in most classic models (PV driven PING). Here, SOM interneurons might provide tonic inhibition that increases the gain of gamma oscillations, but does not contribute significantly to synchronizing the E cells (Börgers and Kopell, 2008); (2) An E-SOM based mechanism (Veit et al., 2017a; Hakim et al., 2018), where SOM neurons are the main synchronizers of the E cells (SOM driven PING); (3) An E-SOM-PV based mechanism, in which both cell types may play specific roles, e.g. by providing inhibition at specific moments in the gamma cycle (Onorato et al., 2023). We note that causal manipulations like silencing a cell type are insufficient to disentangle these scenarios, as a suppressive effect on gamma by silencing SOM cells (Veit et al., 2017a) may be consistent with all three models.

**Figure 1.**
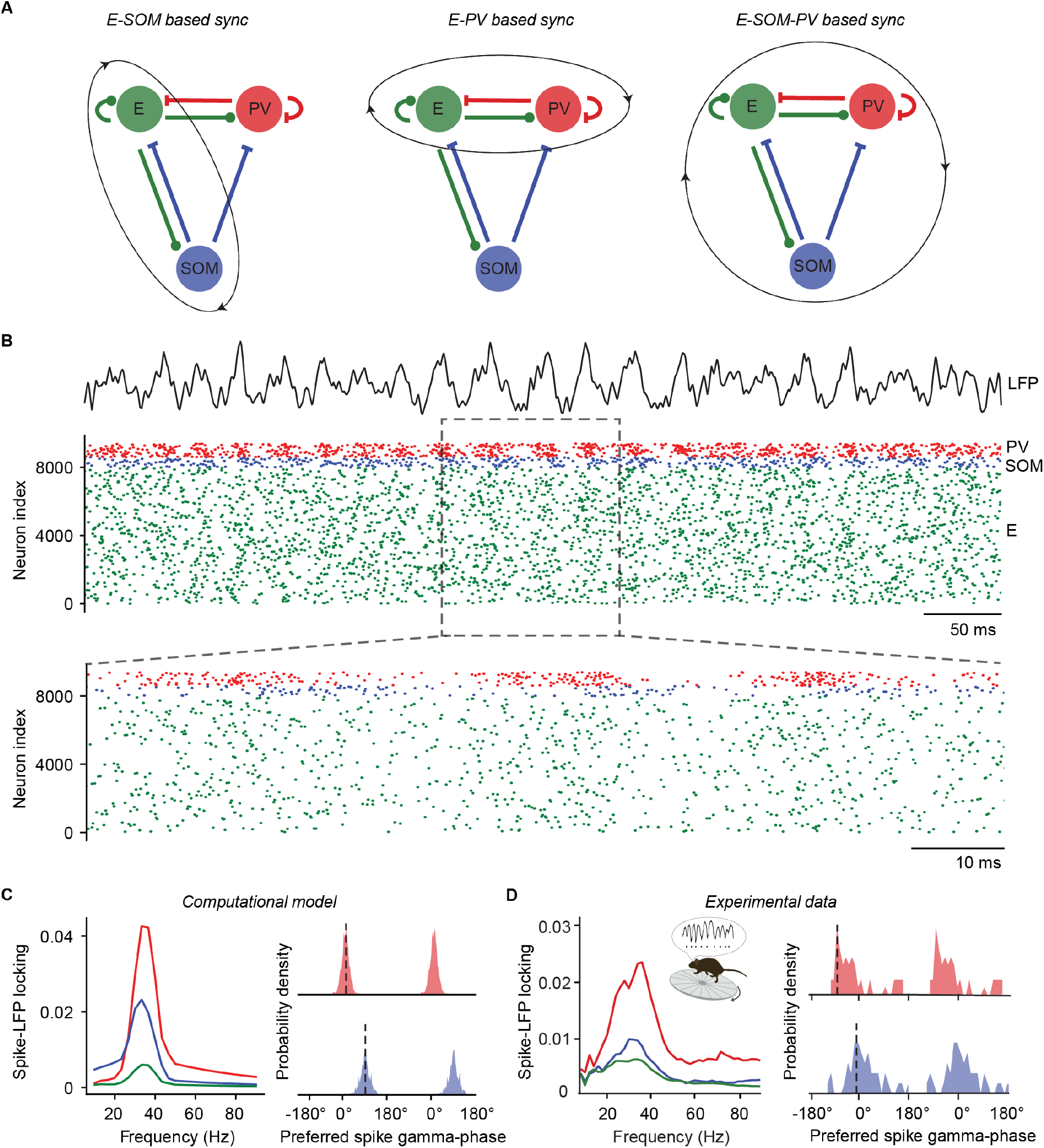
Canonical model reproduces E, PV, and SOM activity during gamma oscillations in V1. **A** Mechanistic hypotheses for gamma generation: E-SOM based synchrony, E-PV based synchrony, and integrated E-PV-SOM circuit based synchrony (left to right). Arrows represent anatomically grounded connections between neural populations **B** Self-organized activity of the canonical E-PV-SOM network. The black continuous line at the top represents the model-generated LFP. In the lower panel we observe the network’s raster plot, where green dots indicate E cell spikes, blue dots represent SOM cell spikes, and red dots show PV cell spikes. A zoomed view of the raster within a shorter time window is displayed at the bottom of the panel. **C** Spike-LFP locking of neuron types (E: green, SOM: blue, PV: red) and phase delay distribution relative to the LFP peak (PV: red, SOM: blue) as predicted by the model. **D** Same as panel **C** for experimental data in mice V1.

Thus, while recent empirical studies have provided important data concerning the dynamics of different cell classes and the effects of causally manipulating them, what is crucially missing is a unified computational understanding of how excitatory, PV and SOM cells interact to generate network oscillations and control network stability. To this end, we built a computational model of three populations (SOM, PV, E cells) constrained by anatomical connectivity data (Pfeffer et al., 2013). We compare the network dynamics and phase-locking of different cell types to those observed *in vivo*, and perform various causal manipulations on the spiking activity of SOM and PV cells to reveal their distinct causal roles. We furthermore perform alterations of the structural composition of the circuit, including connectivity patterns and hierarchical gradients in SOM and PV density, to determine network stability and the characteristics of oscillations.

## Results

### E-PV-SOM canonical microcircuit captures key features of in vivo cortical network dynamics

We constructed a biologically realistic computational model of the cortical microcircuit. The network consisted of 8000 excitatory neurons and 800 inhibitory interneurons representing PV+ cells and 600 inhibitory interneurons representing SOM+ cells (as estimated in V1 (Tremblay et al., 2016)). The connectivity between the cell types was based on the canonical microcircuit described for the mouse visual cortex (Pfeffer et al., 2013). In this microcircuit, the connections of SOM and PV interneurons are distinct: SOM cells project to PV, but not vice versa, and SOM cells have little to no recurrent connections to other SOM cells. In our initial simulations, PV cells and excitatory neurons received a stochastic (homogeneous Poisson) feedforward input, while SOM neurons did not (see Methods). Notice that each E and PV neuron receives an independent Poissonian spike train at the same mean rate.

The simulated network self-generates gamma-band oscillatory activity with a frequency around ∼ 35 Hz, consistent with empirical findings in the visual cortex (Veit et al., 2017a; Onorato et al., 2023; Spyropoulos et al., 2022). Gamma oscillations did not consist of regular periodic cycles but localized events, appearing on top of an overall asynchronous irregular spiking activity (Fig. 1B).

In empirical studies, a common method to characterize the participation of individual neurons in network synchronization is by quantifying the phase-locking of spikes to the ongoing network oscillation (Csicsvari et al., 2003; Hasenstaub et al., 2005; Vinck et al., 2013; Onorato et al., 2020, 2023). To generate a proxy of the LFP signal, we summed the activity of all excitatory neurons together, convolved with a 1ms Gaussian kernel. We then computed spike-LFP phase-locking by using the pairwise phase consistency (PPC) (Vinck et al., 2012), which is a measure unbiased by spike count. Importantly, this measure was used in previous empirical studies, allowing a comparison between the computational model and empirical data. We computed the spike-LFP phase-locking for each individual neuron and then averaged the values within each subpopulation. We found that the phase-locking of PV spikes was higher than that of SOM cells, which in turn was higher than that of E cells (Fig. 1C). The average strength of phase locking as well as the differences between cell types match with a recent empirical study that analyzed the gamma phase-locking of PV, SOM and excitatory neurons in mouse primary visual cortex (Onorato et al., 2023) (Fig. 1D).

Next, we analyzed the phase delays between different cell types. A recent empirical study suggests a substantial phase delay between PV and SOM cells in the gamma cycle, with delayed firing of SOM cells around 6ms (Fig. 1D). Consistent with these empirical observations, we observed a similar lag between the firing of PV and SOM interneurons in the gamma cycle (around 7ms or 80^*°*^).

In sum, an anatomically constrained computational network model closely reproduces the firing dynamics of PV, SOM and excitatory neurons observed in empirical data.

### E-PV-SOM canonical microcircuit captures key features of evoked network activity during visual stimulation

Next, we examined whether this microcircuit recapitulated the stimulus correlates of gamma-band activity, which have been characterized in a large number of previous studies (Veit et al., 2017a; Onorato et al., 2020; Ray and Maunsell, 2010; Uran et al., 2022; Peter et al., 2019a; Vinck and Bosman, 2016).

One key experimental observation is that the V1 gamma-frequency increases with stimulus luminance-contrast (Roberts et al., 2013; Ray and Maunsell, 2010). To model the luminance-contrast dependence, we gradually increased the rate of the external feedforward spike train to E and PV cells and examined the effect on network dynamics. As the intensity of the stimulus increased (see Fig. 2A), the gamma peak-frequency increased, monotonically increasing from ∼ 7 Hz to ∼ 80 Hz. This finding is in agreement with experimental observations (Roberts et al., 2013). The observation of a substantial phase delay between PV and SOM interneurons generalizes across this wide range of input intensities. From lower (low input drive) to higher (high input drive), the time delay decreases from about 10 ms to 4 ms, with a corresponding increase in phase delay. Notably, SOM neuron phase delays and the increase in gamma frequency are also observed during transient increases in feedforward input (bottom panel in Fig. 2A)

**Figure 2.**
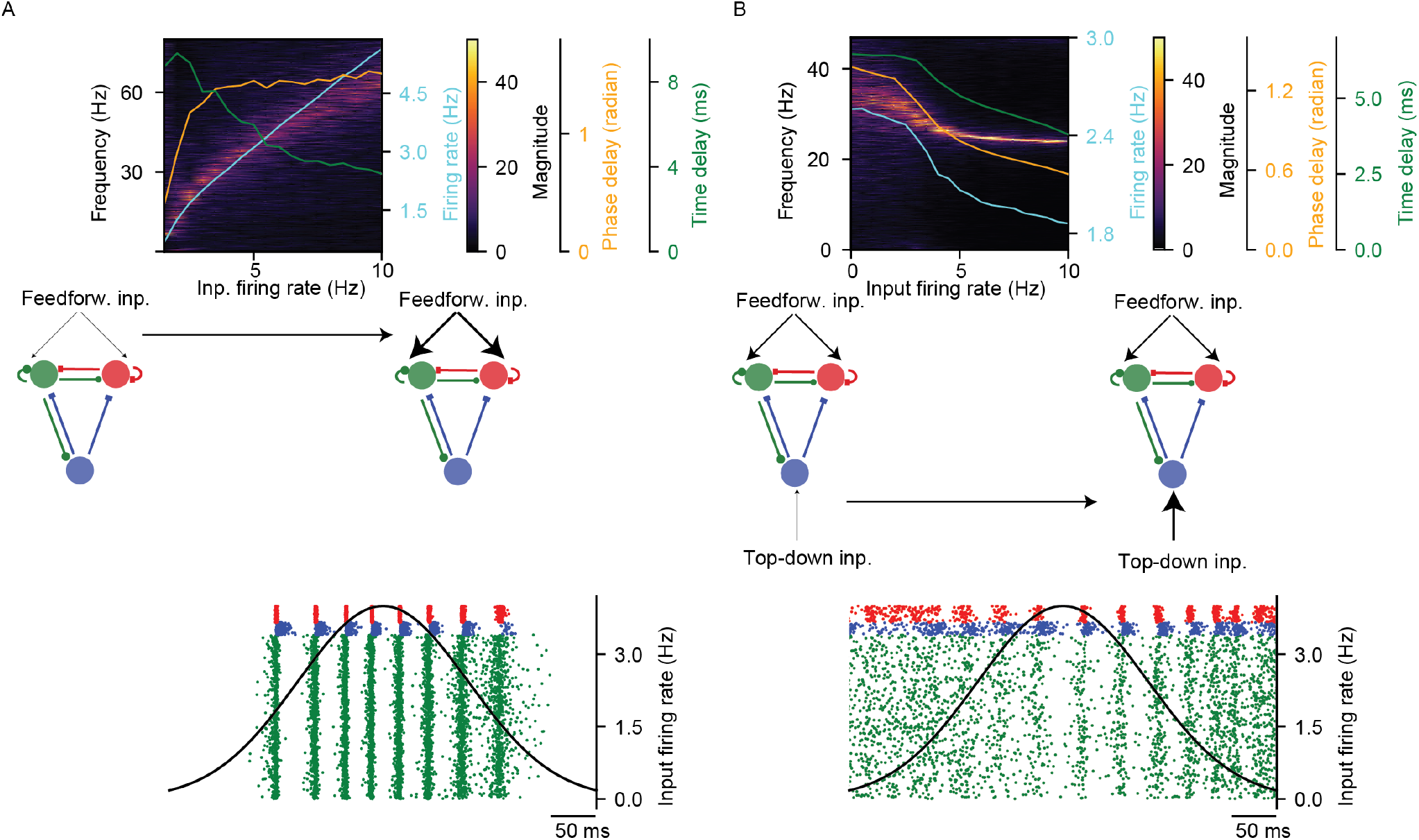
Canonical model reproduces experimental findings on stimulus driven activity in V1. **A** Power spectrum (color-coded) of the LFP at increasing intensity of the tonic feedforward input targeting E and PV cells. Input intensity corresponds to the firing rate of independent Poisson-distributed feedforward spikes received by E and PV cells. The blue line represents the time-averaged E neurons firing rate, while the orange (green) line indicates the mean phase delay between PV and SOM neurons measured in radians (ms). The bottom panel shows the raster plot in response to a transient, time-varying feedforward stimulus (black line superimposed). **B** Same as panel **A**, but with fixed feedforward input (feedforward input firing rate of 4 Hz) and increasing intensity of top-down input targeting only SOM cells.

Another experimental observation is that larger visual stimuli yield stronger gamma oscillations with a lower frequency, which is accompanied by a reduction in firing rates (Gieselmann and Thiele, 2008). This effect is thought to depend on increased activation of SOM interneurons via lateral and top-down feedback (Adesnik et al., 2012). To model this observation, we kept the feedforward stimuli constant (at 4 Hz rate) and introduced a top-down input to the SOM cells (see Fig. 2B). A gradual increase in top-down input to SOM cells led to an increase in the amplitude and a decrease in the frequency of gamma oscillations. These changes in gamma oscillations were accompanied by a reduction in the firing rates of excitatory neurons. The phase lead of SOM over PV interneurons was observed across this range of top-down input drives and also for non-stationary drives.

Thus, the computational model presented here closely recapitulates key experimental observations on the dependence of V1 gamma oscillations on luminance-contrast and surround suppression.

### SOM and PV interneurons suppression

We then used the computational model of the cortical micorcircuit to test for the distinct causal roles of PV and SOM interneurons in governing network dynamics. Previous work has used optogenetic perturbations of both SOM and PV sub-populations to investigate their distinct causal roles in gamma rhythmogenesis. In particular, suppression of SOM interneurons has been shown to reduce the power of gamma band synchronization (Veit et al., 2017a; Chen et al., 2017). By contrast, suppression of PV cells leads to an increased absolute spectral power across a broad frequency range (Veit et al., 2017a; Chen et al., 2017). Interestingly, higher levels of PV neurons suppression have been tested in these experiments, often resulting in uncontrolled network dynamics, evidenced by epileptic like activity (Veit et al., 2017a). It is important to notice that these experiments showed also that a moderate silencing of PV or SOM does not change the frequency of oscillations (Veit et al., 2017b; Chen et al., 2017).

To verify if our model was consistent with these experimental findings, we performed numerical simulations by silencing interneurons from either the PV or SOM populations. in order to mimick the inactivation of intenrneurons, we simply removed a fraction of PV or SOM neurons from the network. The strength of silencing was indexed by the fraction of silenced inhibitory interneurons. Increased suppression of SOM interneurons led to a gradual reduction in gamma-band oscillations, abolishing them with the suppression of most of SOM interneurons (Fig. 3A, C). At the same time, the peak gamma-frequency did not show a substantial change with suppression of SOM interneurons.

**Figure 3.**
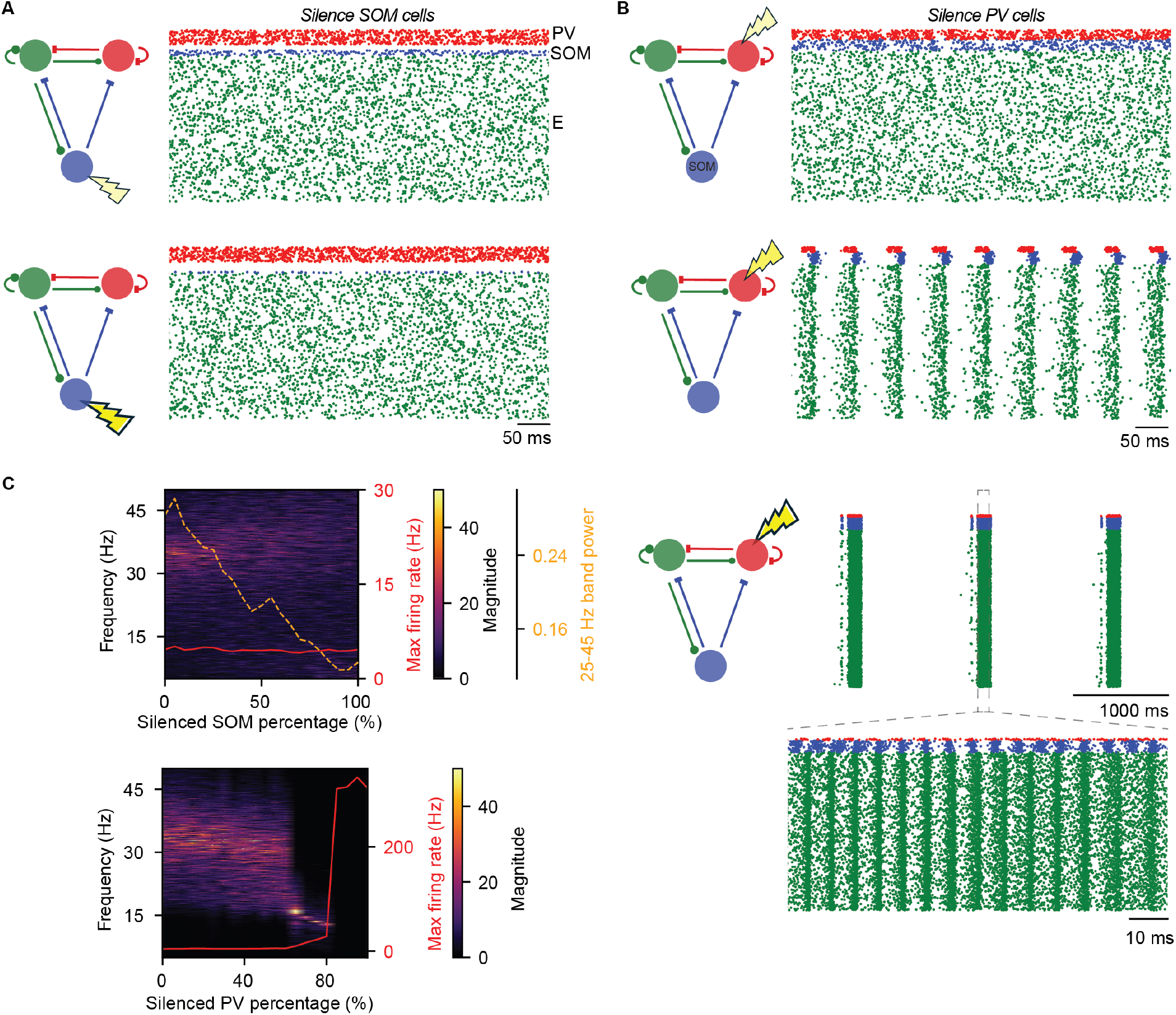
Distinct effect of silencing SOM and PV interneurons on network dynamics. **A** Top and bottom panels display raster plots corresponding to the silencing of 50% and 90% of SOM neurons, respectively. **B** Top, middle and bottm panels display raster plots corresponding to the silencing of 40%, 70% and 90% of PV neurons, respectively. A zoom in the hyper-synchronous epileptic phase is shown at the very bottom. **C** Top panel shows the power spectrum (color-coded) of the LFP at increasing fraction of suppressed SOM cells. Orange dashed line shows the mean power in the band between 25 to 45 Hz. Red continuous line shows the maximum (over time) E cells firing rate. The bottom panel shows the same as the top panel but for an increasing fraction of suppressed PV cells.

Suppression of PV interneurons produced effects that were distinct from silencing SOM interneurons. At lower silencing intensities (20 – 50% stimulation, see top panel in Fig. 3B, C), gamma-amplitude and frequency showed only minor changes. However, beyond 60% PV silencing, a sudden transition to periodic (not transient) synchronous oscillations at lower frequencies (10-15 Hz) is observed at higher level of PV silencing (∼ 60%, see middle panel in Fig. 3B, C). At very strong levels of PV silencing, beyond ∼ 80%, the network exhibited periods of hypersynchronous seizure-like activity. This seizure-like activity was characterized by high-frequency oscillatory activity around ∼ 150% Hz that was periodically interleaved by periods silence, and a strong increase in firing rates of excitatory neurons (bottom panel in Fig. 3B).

These results agree well with previous empirical observations of optogenetic manipulation of SOM and PV neurons Chen et al. (2017); Veit et al. (2017b). Interestingly, they suggest that both intact SOM and PV interneuron activity is necessary for the generation of gamma oscillations, with SOM neurons primarily controlling the amplitude of oscillations and PV interneurons controlling the frequency. Furthermore, PV interneurons control network stability, while SOM interneurons do not.

### Spike-time-disruption of PV and SOM interneurons

The results on the suppression of SOM and PV interneurons suggest causal roles in the genesis of gamma oscillations. However, it is possible that, for example, sufficient activation of SOM (or PV cells) is necessary to provide sufficient inhibitory drive, but that the precise timing of SOM spikes is not necessary to generate gamma oscillations. In other words, the effect of silencing SOM (or PV cells) may merely indicate a role as *modulator*, but not as *pacemaker*, driving the precise timing of excitatory-cell firing. To investigate this, it is necessary to perturb the precise spike timing of both SOM and PV cells, while keeping their firing rates constant. Such a manipulation would be extremely challenging *in vivo*, but is feasible in the presented computational model.

In the computational model, we gradually jittered the timing of spikes by replacing a fraction of neurons with random spike trains generated by a homogeneous Poisson process, while maintaining their original mean firing rate (see Methods). This spike-time-disruption effectively decoupled either part of the SOM or PV interneuron population from the rest of the network while preserving their overall mean modulatory effect. Increased spiketime-disruption of SOM cells led to a gradual decrease in gamma-band amplitude with no major effect on gamma frequency, and no effect on network stability (see Fig. 4A), closely resembling the effects of silencing the SOM cells. Spike-time-disruption of PV interneurons led to a gradual decrease in the oscillation frequency from about 35 to 10 Hz (see Fig. 4B) with an increase in oscillation amplitude. This gradient decrease in oscillations’ frequency was not observed in the previous manipulation of silencing PV cells. Spike-time-disruption of more than ∼ 80% of PV cells led to seizure-like activity, with massively increased firing rates of excitatory neurons.

**Figure 4.**
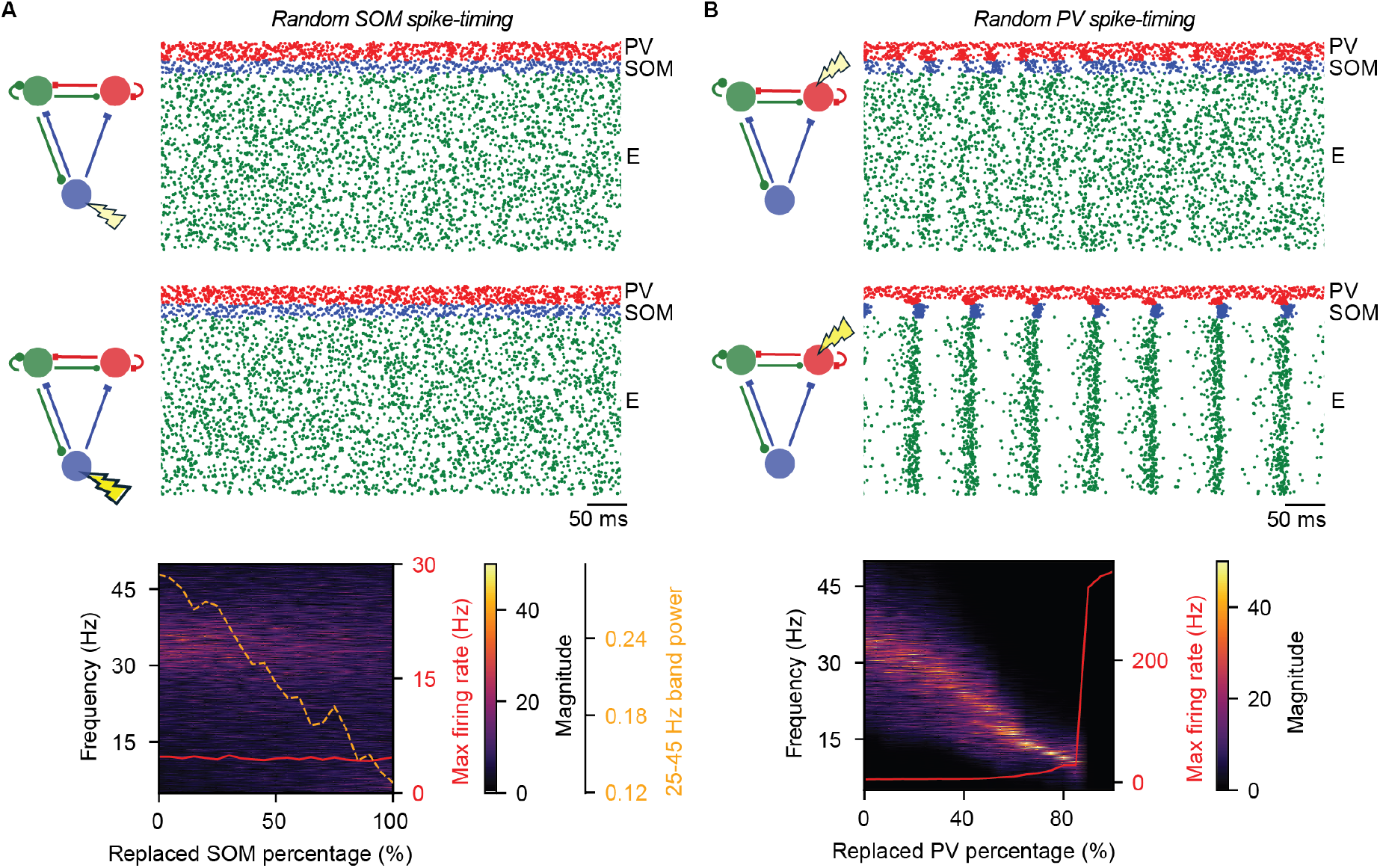
Network effects of randomizing spiking times of SOM and PV interneurons. **A** Top and middle panels display the raster plots corresponding to the replacement of 50% and 90% of SOM neurons (respectively), whose spikes are substituted with a Poissonian spike train at the same mean rate. The bottom panel displays the power spectrum (color-coded) of the LFP at increasing fraction of replaced SOM cells. Orange dashed line shows the mean power in the band between 25 to 45 Hz. Red continuous line shows the maximum E cells firing rate. **B** Same as panel **A** but for the replacement of PV neurons. Top and middle panel correspond to the replacement of 40% and 70% of PV neurons.

The spike-time-disruption manipulation reveals that PV and SOM interneurons do not merely modulate network synchrony, but that the precise spike timing of both cell types is necessary for maintaining gamma oscillations. The specific spike timing of SOM interneurons controls the amplitude of gamma oscillations, while the main effect of PV interneurons is to control the frequency of network oscillations. Furthermore, our findings suggest that the precise spike timing of PV interneurons is causally necessary for network stability, i.e. tonic PV inhibition is unsufficient to yield stable network dynamics.

### Network dynamics is governed by PV-SOM asymmetric connectivity

A prominent feature of the canonical E-PV-SOM microcircuit lies in the asymmetry in the connectivity between different interneurons, defined by the absence of PV-to-SOM coupling and recurrent SOM-to-SOM coupling. We therefore wondered how this asymmetric connectivity profile influences the generation of network synchronization. To this end, we modified the connectivity scheme in our model and performed new numerical simulations.

First, we examined the effect of including mutual inhibitory connections between PV and SOM interneurons, by increasing the inhibitory feedback from PV to SOM interneurons. Increasing inhibitory PV-to-SOM coupling led to a substantial reduced gamma oscillation power and transition towards asynchronous dynamics (Fig. 5A). Gamma oscillations are thus abolished by introducing symmetric (i.e. reciprocal) PV-SOM interactions.

**Figure 5.**
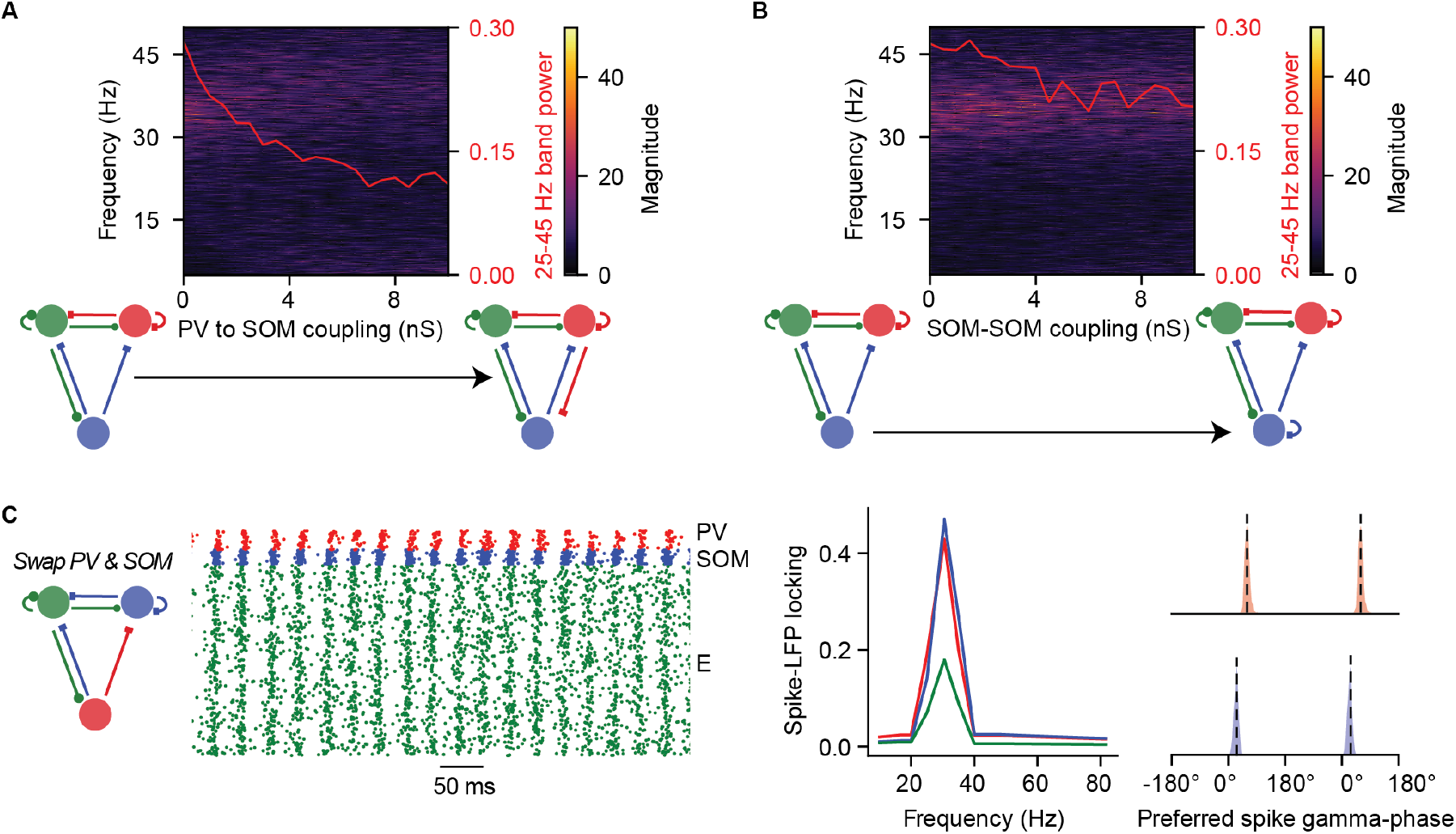
Gamma oscillations arise from PV-SOM asymmetric connectivity. **A** Power spectrum (color-coded) of the LFP at increasing strength of the PV to SOM coupling. Red continuous line shows the mean power in the band between 25 to 45 Hz. **B** Same as **A** but for increasing stength of the SOM to SOM coupling (here the coupling from PV to SOM is set to zero, as in the control microcircuit of Fig. 1). **C** Simulation of a network with swapped PV and SOM neurons, preserving the original connectivity scheme of Fig. 1 (thus here we have no PV-to-PV and no SOM-to-PV connections). We report the raster plot on the left, the Spike-LFP locking in the middle panel and the Preferred spike gamma-phase as for Fig. 1C on the right panel. Notice that red lines (and dots) correspond to PV and blue lines (and dots) to SOM.

**Figure 6.**
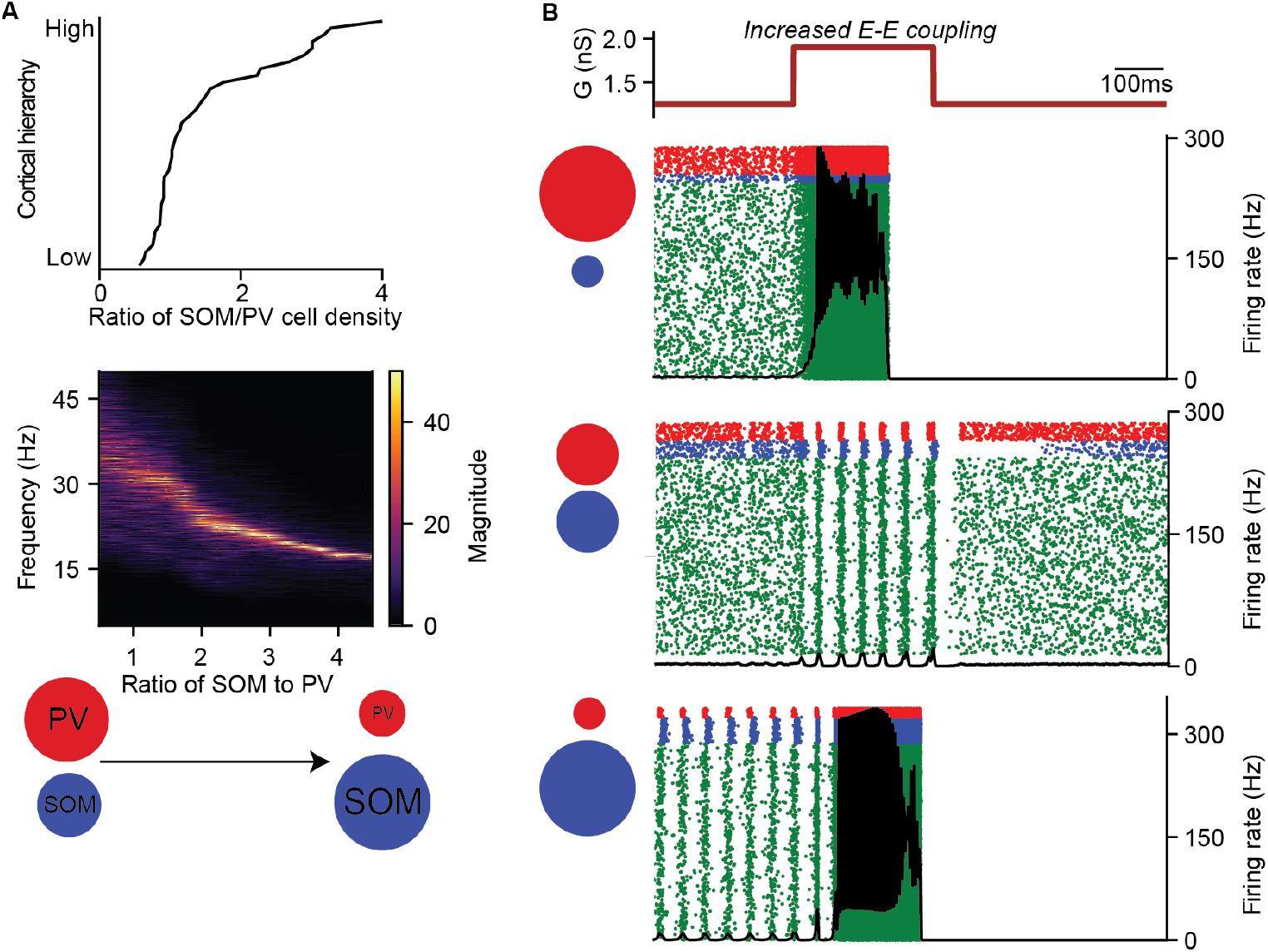
Oscillation frequency and epileptic stability gradients across the cortical hierarchy A. The top panel shows the gradient of the density ratio between SOM and PV cells across the cortical hierarchy (data from Kim et al. (2017)). Lower panel displays the power spectrum (color-coded) of the LFP at the different values of SOM to PV ratio. In these simulations the total amount of interneurons is fixed, the respective number of PV and SOM cells is varied. **B** We conducted numerical simulations on the microcircuit, applying a localized increase in E-to-E coupling strength for a duration of 300 ms (top panel). To examine the impact of interneurons composition, we recorded the corresponding raster plots for networks with different SOM-to-PV ratios: 0.3 (top raster plot), 1 (middle raster plot), and 3 (bottom raster plot). These simulations reveal how variations in interneuron density influence network activity and stability.

Second, we examined the effect of including SOM-to-SOM inhibitory connections. Increasing SOM-to-SOM coupling caused a relatively minor reduction in gamma power (Fig. 5B).

Third, we examined the effect of exchanging SOM and PV interneurons in the microcircuit. This allowed us to examine whether the differences between SOM and PV interneurons in terms of gamma phase-locking and the preferred phase of firing (Figure 1) were due to their intrinsic properties, or rather their position in the E-SOM-PV circuit. In our simulations, SOM and PV interneurons were exchanged while preserving the original connectivity scheme (see Methods). After exchanging the position of SOM and PV interneurons, we now observed a phase advance of SOM interneurons relative to PV cells, i.e. a reversal of the phase-delay between SOM and PV cells as compared to the standard microcircuit. Furthermore, the microcircuit with SOM and PV cells exchanged exhibited relatively strong phase locking to the LFP as compared to the standard network, while the E neurons phase locking remains lower. (Fig. 5C).

These results suggest that the emergent oscillatory dynamics of the network are fundamentally driven by the unique anatomical connectivity scheme between interneurons, in particular the asymmetry between SOM and PV interneurons. Furthermore, these results suggest that the phase delay between SOM and PV interneurons results from their position and connectivity in the microcircuit, rather than from differences in their intrinsic electrophysiological properties or in ad-hoc delays in spike transmission.

### Emergent time scales and network stability across the cortical hierarchy depend on SOM/PV density ratio

Building on our model of the primary visual cortex (V1), we extended our analysis to explore model predictions beyond V1. The V1 computational model contained slightly more PV than SOM interneurons (Tremblay et al., 2016). However, experimental studies Kim et al. (2017) report a strong gradient in the SOM-to-PV density ratio across the cortical hierarchy (see Fig. 6A). That is, higher areas (like prefrontal cortex, parahippocampal areas) are characterized by relatively high fractions of SOM cells whereas lower areas (like sensory and primary motor areas) are characterized by relatively low fractions of PV cells. Another experimental observation is a systematic increase in intrinsic time-scales and decrease in oscillation frequencies across the cortical hierarchy (Murray et al., 2014; Gao et al., 2020; Vinck et al., 2023). We thus wondered, how an increase in the SOM-to-PV ratio changes the network dynamics, and whether increased SOM-to-PV ratios entail slower network dynamics.

To investigate this question, we performed numerical simulations by systematically adjusted SOM-to-PV ratios, while keeping the total number of interneurons and input drive fixed. With increasing SOM-to-PV ratios, we observed a gradient of network oscillation frequency decreasing from gamma frequencies (∼40 Hz) towards slower and higher-amplitude alpha/beta oscillations (∼15 Hz). That is, a higher SOM-to-PV ratio indeed predicts slower network dynamics, consistent with empirical observations.

We wondered whether the SOM-to-PV ratio has implications for the stability of the network, which can be examined by increasing the recurrent excitatory (E-E) connectivity of the network. We performed numerical simulations by introducing a transient local increase in excitatory-to-excitatory conductances (top panel, Fig. 6B). Our analysis shows that low SOM to PV density ratios (*SOM/PV <* 0.4) and high SOM to PV density ratios (*SOM/PV >* 2.5) are prone to destabilization with increased recurrent excitation, leading to seizure-like activity characterized by high firing rates. In contrast, networks with intermediate levels of SOM to PV ratios (0.4 *< SOM/PV <* 2.5) did not stabilize despite increasing recurrent excitation, and instead exhibited gamma/beta network oscillations (Fig. 6B). These findings indicate that the SOM-E-PV microcircuit with intermediate ratios of SOM and PV interneurons exhibits enhanced stability as compared to circuits that predominantly contain either SOM or PV interneurons.

## Discussion

Recent experimental studies used optogenetic techniques to characterize the phase-locking of distinct GABAergic interneurons and the effects of silencing them (Onorato et al., 2023; Veit et al., 2017a; Chen et al., 2017). Yet a comprehensive computational understanding of their distinct causal roles for cortical dynamics remains to be established. To this end, we built a computational model of three populations (SOM, PV, E cells) constrained by anatomical connectivity data (Pfeffer et al., 2013). Our model reproduces key experimental findings, including: stochastic gamma oscillations (Spyropoulos et al., 2022; Burns et al., 2011); the distinct phase-locking and timing of PV and SOM interneurons in the gamma cycle (Onorato et al., 2023); the dependence of gamma on feedforward and feedback inputs (Ray and Maunsell, 2010; Gieselmann and Thiele, 2008; Veit et al., 2017a; Jadi and Sejnowski, 2014; Roberts et al., 2013); and the effect of optogenetic perturbation of SOM and PV cells (Veit et al., 2017a; Chen et al., 2017). Besides these findings, we performed various causal manipulations yielding novel predictions: First, we show that the precise spike timing of both PV and SOM interneurons is necessary for gamma oscillations’ generation, with distinct functional roles for PV and SOM interneurons: PV cells regulate oscillation frequency and network stability, whereas SOM cells primarily control the oscillation amplitude. Second, we show that the asymmetric connectivity between SOM and PV is a key mechanism for generating the delayed firing of SOM relative to PV neurons, which is necessary for the generation of gamma oscillations. Third, the gradient in SOM/PV interneurons across the cortical hierarchy (Wang, 2020) predicts a decreasing gradient in cortical oscillation frequencies, consistent with experimental data (Murray et al., 2014; Gao et al., 2020; Vinck et al., 2023). Specifically, a heterogeneous, balanced mixture of SOM/PV interneurons produces stochastic gamma oscillations with a relatively low propensity for generating seizures. By contrast, a high density of either SOM or PV cells produces, respectively, low-frequency oscillations or asynchronous activity, with an elevated seizure propensity. Thus, the three-population E-SOM-PV circuit appears to confer functional benefits in terms of network stability compared to two-population E-SOM or E-PV circuits. We refer to the computational mechanism for network oscillations, which is distinct from the classic E/I models such as the PING model, as the CAMINOS model (CAnonical MIcrocircuit Network Oscillations).

### Prediction of experimental findings

The CAMINOS model described here reproduces a range of experimental findings, which include: (1) The stochastic gamma oscillations as observed in visual cortex (Spyropoulos et al., 2022; Burns et al., 2011), which can be described as quasi-oscillations (Wallace et al., 2011). (2) The model provides a very close fit to the optogenetic results of Veit et al. (2017a), with similar results for silencing SOM and PV interneurons, such as the sudden phase transition for silencing PV interneurons and a gradual decrease in gamma amplitude for silencing SOM interneurons. We predict that experimentally, there should be a sudden transition from gamma to low frequency oscillations to seizure activity when silencing PV interneurons, which was likely not observed in Veit et al. (2017a) as only a limited range of light intensities was used to silence PV interneurons. Chen et al. (2017) did observe a decrease in gamma oscillations together with a shift towards lower frequencies with PV silencing, which matches the predictions of the CAMINOS model. It is possible that the difference between Chen et al. (2017) and Veit et al. (2017a) reflects stronger silencing PV interneurons in Chen et al. (2017). (3) We reproduce the experimentally observed phase delay between PV and SOM interneurons (Onorato et al., 2023). We show that this phase delay is robust across different input frequencies, suggesting that the delay is a property that generalizes beyond gamma oscillations. Indeed, Onorato et al. (2023) observed that a delayed firing of SOM relative to PV interneurons was also observed w.r.t. the transient evoked by a stimulus onset. (4) The model reproduces the characteristic increase in frequencies with an increase in input drive, which has been observed by manipulating stimulus luminance-contrast in primary visual cortex (Ray and Maunsell, 2010; Roberts et al., 2013). Furthermore, the model can account for the observation that large visual stimuli cause a decrease in gamma frequency and increase in gamma amplitude, accompanied by decreased firing rates. In our model, the effect of increasing stimulus size was modelled by an increase in feedback drive onto SOM interneurons. Indeed, studies have shown that large and spatially predictable stimuli, which induce strong gamma-band activity at a relatively low gamma frequency (Uran et al., 2022; Gieselmann and Thiele, 2022; Vinck and Bosman, 2016; Peter et al., 2019b), evoke strong activation of SOM interneurons (Adesnik et al., 2012; Keller et al., 2020). (5) The CAMINOS model predicts that cortical circuits with a higher density of SOM than PV cells produce oscillatory dynamics at lower frequencies, i.e. in the alpha/beta range. This prediction agrees with the idea that network dynamics are faster in lower hierarchical levels of the cortex and decrease in frequency across the hierarchy (Murray et al., 2014; Gao et al., 2020; Vinck et al., 2023), at least in the activated cortical state. For instance, gamma dynamics in the activated state are frequently observed primary visual cortex, somatosensory cortex or olfactory bulb, while beta-band activity is typically observed in intermediate areas of the hierarchy during active states (Vinck et al., 2023). From this perspective, different cortical areas may generate oscillations through the same microcircuit interactions, which are expressed at different frequencies because of varying SOM/PV densities across the cortical hierarchy. Importantly, in the CAMINOS model, the network frequency also increases as a function of cortical activation. Thus, the CAMINOS model predicts that a hierarchy of time constants can only be observed if the areas are activated by sensory inputs or actively engaged in a task. This drive dependence may explain why, in a given behavioral task, large parts of the primary visual cortex may engage in low-frequency activity, with gamma-activity being sparsely expressed (Hoffman et al., 2024; Vinck et al., 2023).

### CAMINOS: canonical microcircuit network oscillations

The CAMINOS mechanism introduced here is distinct from the classic PING or Wilson-Cowan model, where oscillations emerge from synaptic delays or differences in intrinsic neuronal time scales between E and I neurons (Wallace et al., 2011; Wilson and Cowan, 1972; Buzsáki and Wang, 2012; Kopell et al., 2000). In CAMINOS, by contrast, oscillations emerge from the asymmetric connectivity between SOM and PV interneurons, which imposes precise and delayed spiking times between distinct interneuron populations that are necessary to sustain gamma oscillations.

In each oscillatory cycle, an increase in excitation first gives rise to a very rapid activation of PV interneurons. However, this rapid inhibitory feedback is itself not capable of sustaining stable oscillatory activity (Figure S1). SOMs activate about 6-7ms later, and this delayed inhibition stabilizes the network oscillation. Yet, by itself the E-SOM interactions support low-frequency rather than high frequency oscillations. Thus, the oscillatory behavior arises from a fast component mediated by PV interneurons and the delayed, inhibitory component mediated by SOM interneurons. This delayed inhibitory SOM activity results from the asymmetric connectivity between PV and SOM interneurons in the microcircuit. This architecture is based on experimental observations (Pfeffer et al., 2013) and provides a distinct mechanistic explanation for oscillations grounded on the heterogeneity of interneurons.

Importantly, the CAMINOS model predicts the wide range of dynamics across the neocortical hierarchy, as well as the changes in frequency within areas as a function of input drive. In classical E/I models, the frequency of oscillations is determined by fixed intrinsic synaptic time constants (Brunel, 2000; Wilson and Cowan, 1972; Nandi et al., 2024). Accordingly, CAMINOS proposes a unifying mechanism for oscillations across different frequencies, driven by network architecture rather than distinct circuit mechanisms or arbitrary gradients in synaptic time scales. From this perspective, oscillations at different frequencies may not inherently correspond to distinct functional roles, but rather reflect gradients between cortical circuits and differences in the strength of feedforward and feedback drive into a circuit.

Thus, the precise dynamics exhibited by the same microcircuit may be diverse and reflect the structure of the feedforward and feedback input drive. If the drive into the network increases, the network may generate a burst of oscillatory activity. Different microcircuits may express these oscillatory bursts at different frequencies, depending on the specific composition of the microcircuit. Such a mechanism may account for the state-dependent occurrence of oscillatory bursts at different frequencies during a given task (Lundqvist et al., 2016; Douchamps et al., 2024). Yet, when the network is activated by a sudden transient, the network may generate an aperiodic transient rather than oscillatory activity (Vinck et al., 2023). From this perspective, visual cortex’s may generate low-frequency oscillations in the alpha frequency range in the absence of input drive; a visual stimulus may then produce a rapid aperiodic transient activation driven by the thalamic feedforward input, which disrupts this oscillatory pattern; this transient is then followed later in time by oscillatory activity in the gamma frequency range given a relatively strong feedforward drive that only varies slowly with time.

### Implications for regulation of network stability

The classic modeling work of van Vreeswijk and Sompolinsky (1996) shows that E/I balance is a prerequisite for network stability, and that strong inhibitory feedback allows for strong recurrent excitatory connections. Here we make several core findings concerning the way in which a three-population E-SOM-PV circuit controls stability:

1. Network stability is enhanced by a network configuration with both PV and SOM interneurons that have a specific asymmetric connectivity profile, as compared to a network that is dominated either by PV or SOM interneurons. One possible mechanism may be the fact that PV and SOM interneurons provide inhibition at different delays, leading to a fast and slow inhibitory feedback. Our causal manipulations show indeed that network stability and excitation/inhibition balance is enhanced by precisely timed spiking of interneurons, which ultimately drive network oscillations. This finding contrasts with classical E-I network models, where oscillations are typically not sustained in the thermodynamic E/I balanced limit (Di Volo et al., 2022). Instead, the CAMINOS model introduced here suggests that oscillations can emerge as a natural and stable solution in balanced networks, governed by the asymmetric and precisely timed interactions between distinct interneuron populations.
2. Interestingly, we show that the network can approach a destabilized, seizure-like state both from the highly synchronized state, in case of a prevalence of SOM interneurons, and the asynchronous state, in case of a strong prevalence of PV interneurons. This predicts that cortical populations may generate two kinds of seizures. The idea that network stability is enhanced by a more heterogeneous circuit fits in principle well with theories that emphasize epilepsy may result from a loss in heterogeneity (Rich et al., 2022). Our findings may suggest that frontal epilepsy or medial temporal lobe epilepsy may potentially reflect enhanced suspectability to seizures due to the specific SOM/PV ratios in those areas, especially when combined with strong recurrent excitatory activity (Elston et al., 2011).
3. The presented computational model further suggests that, in a network with mixed PV and SOM cells, PV cells may be the main stabilizers of network activity, consistent with the empirical observation that optogenetic silencing of PV interneurons leads to network instability (Veit et al., 2017a). However, SOM neurons do contribute to stability, but we predict that a strong perturbation, such a increasing recurrent excitatory connectivity, is necessarily to reveal this dependence in circuits with a high density of PV interneurons.
4. Surprisingly, we show that the precise timing of PV spiking matters for stability, even on fast time-scales. Hence, a decoupling of PV with network activity may facilitate the emergence of seizure activity. Indeed, we have previously shown that a decoupling of PV cells, but not SOM cells, with ongoing network activity in the gamma-frequency range was observed preceding pharmacologically induced seizures in hippocampus CA1 (Miri et al., 2018).

Future work should address the host of experimental predictions made by the CAMINO model, e.g. by perturbing the timing of SOM and PV firing, modulating the connectivity between SOM and PV interneurons, and perturbing SOM and PV interneurons across the hierarchy. Another important open question is how the precise, structured pattern of feedforward drive in a network with non-random connectivity weights modulates gamma-band activity, in order to test its potential functional role in predictive coding and stimulus inference (Uran et al., 2022; Vinck et al., 2023; Echeveste et al., 2020).

## Methods

### Neurons’ model

For simulating the dynamics of neurons’ membrane potential *V*(*t*) we used the Adaptive Exponential integrate and fire model (AdEx) described by the following differential equations Brette and Gerstner (2005):

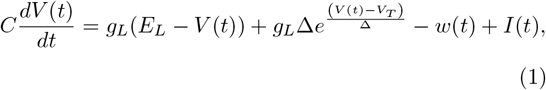

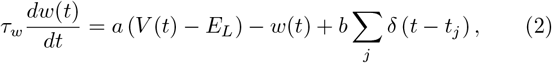

where *t* represents time, *C* is the neuron membrane capacitance, *g*_*L*_ is the leakage conductance, *E*_*L*_ is the resting membrane potential, *V*_*T*_ is the effective threshold, and Δ is the threshold slope factor, determining the width of the action potential. The term *I*(t) represents the sum of all currents injected into the neuron due to the spiking activity of presynaptic neurons. The variable *w*(*t*) is represents an adaptation current, a sort of activity-dependent self-inhibition (see the negative term −*w*(*t*) in Eq. (1)). Whenever the neuron emits a spike at time *t*_*j*_, the adaptation avriable *w* is increased of an amount *b* because of the integration of the Dirac function *δ*(*t* − *t*_*j*_). The parameter *a* represents the subthreshold adaptation, which depends instead by the dynamics of *V* (*t*). The variable *τ*_*w*_ is the adaptation time scale of decay. The *j* − *th* spiking time *t*_*j*_ is defined as the time at which the membrane potential reaches the threshold, defiend as *V*_*T*_ + 5Δ. After the spike generation, the membrane potential is reset to *V*_reset_ at time 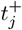, and kept fixed for a refractory period *T*_ref_. Neuron parameter values using for simulations are gathered in Table 1. These parameters have been chosen according to electrophysiological insights of E, SOM and PV neurons. The width Δ of the action potential is larger in SOM and E neurons with respect to PV (McCormick et al., 1985; Hu and Jonas, 2014; Casale et al., 2015). Moreover, E and SOM neurons are known to be characterised by spike frequency adaptation with respect to PV (McCormick et al., 1985; Kawaguchi and Kubota, 1996; McGarry et al., 2010; Tremblay et al., 2016). The different values of *E*_*L*_, setting neurons’ excitability, across the three neurons are tuned in order to quantitatively fit the mean firing rate of neurons in V1, with higher firing rates for PV and lower rates for SOM and RS (Ma et al., 2010). The rest of the values are identical between different neurons’ types and are tuned to their standard values in biologically realistic range.

**Table 1:** Neurons’ parameters

| Parameter | E, PV, SOM |
| --- | --- |
| $V_T$ | -50 mV |
| $\Delta$ | 2, 0.5, 1.5 mV |
| $T_{\text{ref}}$ | 5 ms |
| $\tau_w$ | 500 ms |
| $a$ | 4, 0, 4 nS |
| $b$ | 130, 0, 25 pA |
| $C$ | 200 pF |
| $g_L$ | 10 nS |
| $E_L$ | -60, -60, -55 mV |
| $V_{\text{reset}}$ | -65 mV |

**Table 2:**
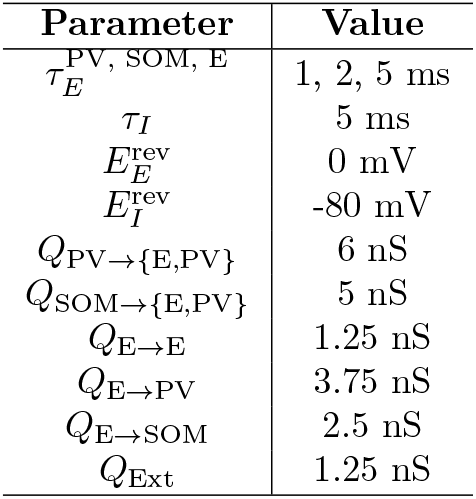
Synaptic parameters

### Synapses and currents

We consider conductance-based synapses, whose postsynaptic current is given by

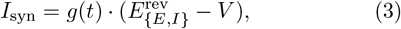

where *V* is the postsynaptic membrane potential and 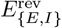 is the reversal potential of the Excitatory (E) or inhibitory (I) synapses. Synapses are classified as excitatory or inhibitory based on the presynaptic neuron type: excitatory (E) or inhibitory (PV or SOM), respectively. The time evolution of the synaptic conductance *g*(*t*) is governed by the following differential equation, which describes both its increase due to presynaptic spikes and its decay over time:

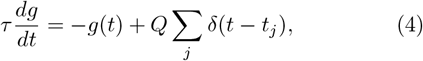

where *δ*(*t* − *t*_*j*_) is the Dirac delta function representing a presynaptic spike at time *t*_*j*_. The parameter *τ* is the time constant of synaptic decay (we use the symbol 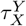 for the synapse from a neuron of the type X to a neuron of the type Y, with {*X, Y*} = *E, SOM* or *PV*). The summation means that at each time a presynaptic neuron spikes at time *t* = *t*_*j*_, the synaptic conductance *g*(*t*) is increased by a discrete amount *Q*, which represents the quantal conductance of the synapse (we use the symbol *Q*_X→Y_ for the synapse from a neuron of the type X to a neuron of the type Y), i.e. the strength and the efficacy of the synapse. Synaptic parameters values are chosen based on electrophysiological proprieties following (Destexhe et al., 1998). Little variation in parameter’s values between synapse type (E to SOM or E to PV) are due to quantitative fine tuning of model’s activity to experimental data (in particular to quantitatively match spike to LFP locking, see Fig.1). The net input current to a neuron, *I*(*t*) (see Eq.(1)), is then the linear sum of all the presynaptic excitatory and inhibitory synaptic currents.

Let us note that we deliberately do not consider any synaptic delay in our model, in order to demonstrate that synaptic delay is not necessary for the genesis of oscillations and that the phase delay between PV and SOM cells spiking times is not due to an ad-hoc choice of synaptic delay. Nevertheless, adding synaptic delays does not affect the overall qualitataive features of the model.

### Number of neurons and network connectivity

We considered *N*_*E*_ = 8000 excitatory (E) cells, *N*_*I*_ = 1400 inhibitory cells, with *N*_*SOM*_ SOM cells and *N*_*P V*_ PV cells such that *N*_*I*_ = *N*_*SOM*_ +*N*_*P V*_. In the canonical model of Fig. 1, we consider *N*_*P V*_ = 800 and *N*_*SOM*_ = 600, i.e. a SOM to PV ration of *N*_*SOM*_ */N*_*P V*_ = 0.75 as estimated in V1 (Tremblay et al., 2016; Campagnola et al., 2022).

The connectivity between E, SOM and PV cells are tuned based on anatomical data (Pfeffer et al., 2013; Campagnola et al., 2022). In particular, based on these data collected in the primary visual cortex PV do not project to SOM, and SOM cells do not inhibit themselves (see Fig. 1A). In order to model such configuration we condsidered, apart from Fig.5, *Q*_*P V* →*SOM*_ and *Q*_*SOM*→*SOM*_ equal to zero. We employed sparse connectivity such that each pre-post synaptic neurons are connected to each other with a probability *p* = 1.7%. Finally, we considered higher recurrent connectivity from E to PV cells (probability of connection 2*p*, still in biologically realistic range) in order to quantitatively fit the higher Spike to LFP locking of PV neurons.

### Numerical implementation of the network model

Network models were simulated using the Brian 2 Python package (Stimberg et al., 2019). Differential equations were numerically integrated using Euler’s method choosing *dt* = 0.1*ms* as the integration time step. We discard the first 2s transient from numerical simulations that may depend on initial conditions.

### External input

External inputs were modeled as arising from external excitatory cells. We consider that the external input arises from Poissonian spikes of *p* ∗ *N*_*E*_ presynaptic excitatory neurons at a fixed firing rate, with the quantal conductance of *Q*_*Ext*_. We consider a feedforward input targeting E and PV cells at a rate 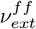 (typically 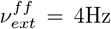 if not stated otherwise). In Fig. 2B we considered also top-down input targeting SOM neurons at a rate 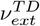 (in allother simulations otherwise 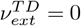).

The input rate 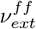 is stationary in all our simulations, apart from data reported in Fig. 2 (bottom panels) where we consider a non-stationary stimuli. In this case, for both 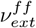 and 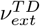, we consider the Gaussian evolution 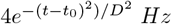, where *t*_0_ is the non-stationary stimulus peak time and *D* = 140ms represents its duration.

### Population activities and LFP

We determine the population activity of one cell type as

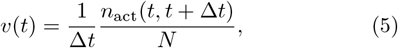

where *N* represents the number of neurons within the population. The term *n*_*act*_(*t, t* + Δ*t*) signifies the cumulative spike count across all neurons in the group within the time window (*t, t* +Δ*t*). Throughout our study, we consistently set Δ*t* to 0.1 *ms* for calculating population activities. After calculating population activities, we convolve *v*(*t*) with a Gaussian kernel of a duration 1ms. In order to generate a proxy for Local Field Potential (LFP), we employ the population activity of E neurons. We remove the first two seconds (transients) of the signal and consider a 10-second duration of LFP signal for our analysis and calculations.

### Frequency analysis and color maps

We performed time frequency analyses on the LFP *x*(*t*). We first removed its dc component, i.e. *x*(*t*) − *x*(*t*), then we normalize it 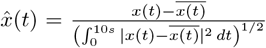. Afterwards, we take its Fourier transform that results in 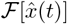 and then calculate the magnitude, 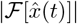. This magnitude is the quantity we report on the color maps in the Figures of the main paper. For some of the color maps, e.g. those of PV manipulations in Fig. 3C, we also perform a bandpass filtering on the signals to prevent showing harmonics on the color maps. In order to estimate the gamma power, we calculated the 25-45 Hz band power via the following quantity 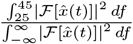.

### Pairwise Phase Locking (PPC)

To quantify the phase locking of spikes to the LFP signal, we use pairwise phase-consistency measure, PPC, (Vinck et al., 2010). We apply this measure on 10 s duration signals. We refer the reader to the mentioned references for calculation details.

### Measuring Spike phase differences

The relationship between spikes and the LFP phase is analyzed by calculating the instantaneous LFP phase at the time of neuronal spikes. We filter the LFP signal around its peak frequency and then we calculate its Hilbert transform from which the instantaneous phase is extracted. For each neuron, the instantaneous phases of the LFP at spike times are recorded. Then to calculate the mean phase of each neuron group we take the average over spike times and all neurons of the group.

### Causal manipulations: Silencing and Desynchronizing

We performed two manipulations on our networks. The first one is silencing, that we simply performed by removing a specific fraction of neurons from the network. The second one is randomizing neurons’ spiking times of a certain fraction of neurons. In practice, we replaced those neurons with with identical independently distributed Poissonian spike trains with a rate equal to the mean rate of neurons before replacing.

### PV and SOM neurons swap

In Fig. 5C, we examined the effects of swapping PV and SOM cells while preserving the original connectivity scheme. In practice, all parameters of the model stay the same, apart from the neoronal parameters of SOM and PV cells (see Table 1) that were exchanged. Moreover, in order to allow the network to be active (PV neurons are silent otherwise) we increased the input rate to PV cells (which is zero otherwise in this new configuration) to 4Hz.

## Acknowledgements

FT and MdV were supported by the French Ministry of Higher Education (Ministére de l’Enseignement Supérieur) and the project LABEX CORTEX (ANR-11-LABX-0042) of Université Claude Bernard Lyon 1 operated by the ANR. MV was supported by an ERC starting grant (850861) SPATEMP, DFG VI Grants (908/5-1 and 908/7-1), an NWO VIDI Grant, and the Dutch Brain Interface Initiative (DBI2).

---

**Figure S1:**
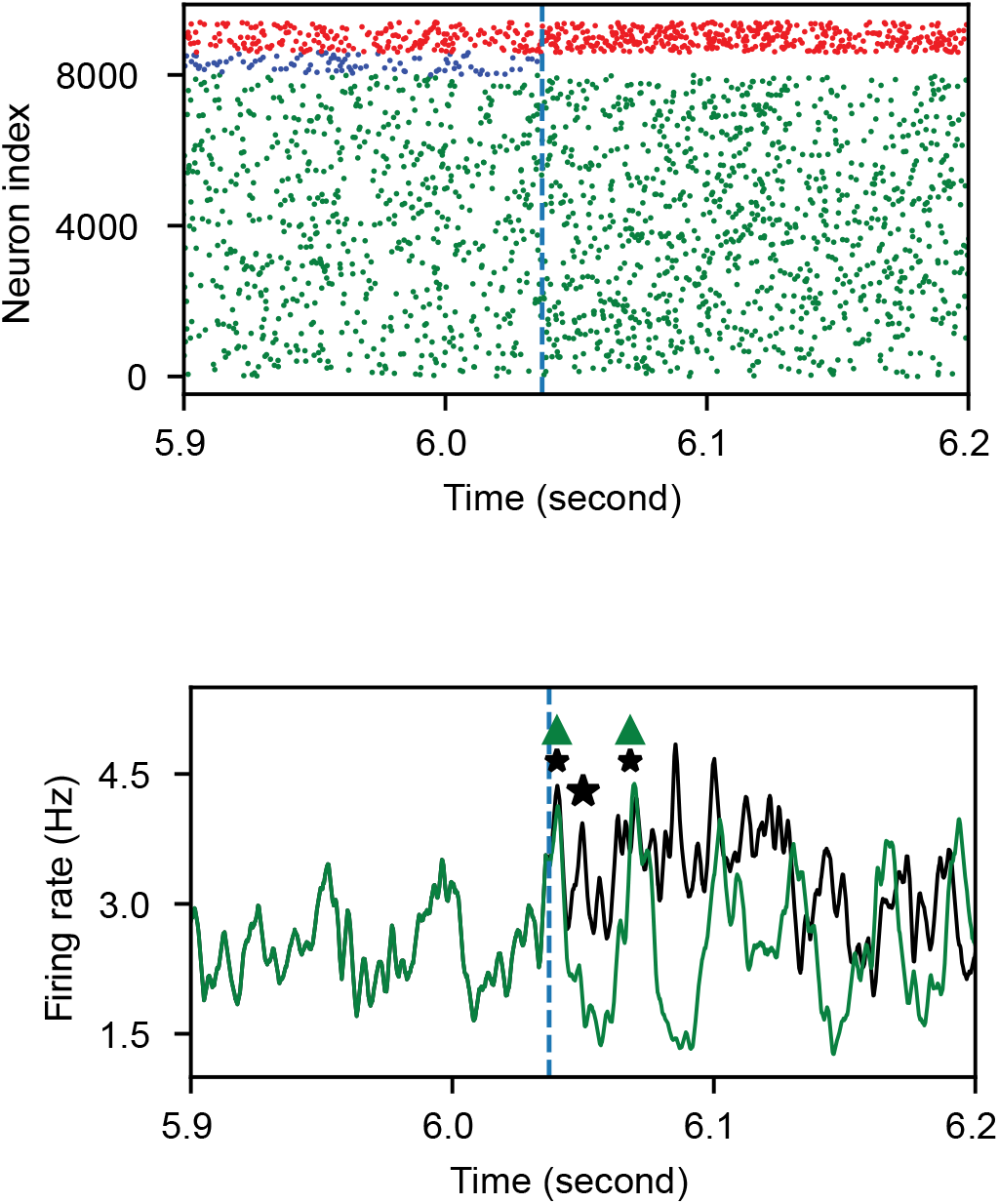
Dynamics of the network in response to silencing SOM activity. In the top panel we report the raster plot, where SOM interneurons are silenced at time ∼ 6.04 s (see vertical blue dotted line). In the bottom panel we report in black (green) the pyramidal neurons firing activity with (without) SOM silencing. After silencing SOM interneurons, the network initially follows the same trajectory (black vs. green line). However, with intact SOM activity, there is a prolonged inhibition that leads to a global deactivation of the network at a gamma frequency. With silenced SOM activity, the network activity increases again after a short interval. However the network is not able to sustain highfrequency oscillations driven by the PV-E loop and transitions to the asynchronous regime.

## Notes

### Competing Interest Statement

The authors have declared no competing interest.

